# BugSeq 16S: NanoCLUST with Improved Consensus Sequence Classification

**DOI:** 10.1101/2021.03.16.434153

**Authors:** Ana Jung, Samuel D Chorlton

**Author notes:** Corresponding Author: Samuel D Chorlton.

## Abstract

NanoCLUST has enabled species-level taxonomic classification from noisy nanopore 16S sequencing data for BugSeq’s users and the broader nanopore sequencing community. We noticed a high misclassification rate of NanoCLUST-derived consensus 16S sequences due to its use of BLAST top hit taxonomy assignment. We replaced the consensus sequence classifier of NanoCLUST with QIIME2’s VSEARCH-based classifier to enable greater accuracy. We use mock microbial community and clinical 16S sequencing data to show that this replacement results in significantly improved nanopore 16S accuracy (over 5% recall and 19% precision), and make this new tool (BugSeq 16S) freely available for academic use at BugSeq.com/free.

---

We read with great interest the recent publication of NanoCLUST, a species-level analysis of 16S rRNA nanopore sequencing data (Rodríguez-Pérez *et al*., 2020). NanoCLUST brings a novel, unsupervised read clustering approach to identify groups of similar reads, and corrects the errors within each cluster to identify a highly accurate amplicon sequence variant (ASV). This approach reduces the typically high (>5%) nanopore sequencing error rate, enabling species-level identification from nanopore 16S sequencing data.

After its release, we rapidly integrated NanoCLUST into BugSeq, our democratized online bioinformatics platform, to provide leading accuracy in 16S nanopore analysis for the academic community, free of charge (Fan *et al*., 2020). While BugSeq users have reaped significant benefits from the development and online deployment of NanoCLUST, we have also noted significant differences from its published accuracy on real world data. Here we report on a frequent discrepancy between NanoCLUST and ground truth data, our solution, and a benchmark of our new classification accuracy.

When designing NanoCLUST, Rodríguez-Pérez et al. used BLAST to classify ASVs (Camacho *et al*., 2009). Specifically, corrected consensus sequences from clusters are BLASTed against the NCBI 16S BLAST database, and the sequence is assigned to the taxonomy of the top hit. This approach leads to a high misclassification rate for several reasons. First, some bacteria have nearly identical 16S sequences, such that only a 100% sequence identity between query and reference sequence is appropriate to assign a sequence to the reference’s species (Edgar, 2018). No such safeguards are taken with the NanoCLUST approach - an ASV may have 80% sequence identity to the top BLAST hit yet be assigned to that species. Second, NanoCLUST accepts a BLAST hit of any length, as long as the expectation value of the alignment is less than 11; that is, for any ASV classification, there may be up to 11 hits found just by chance.

We sought to replace the ASV classification mechanism of NanoCLUST with a more accurate 16S classifier. We tried IDTAXA and a naive Bayes classifier, but settled on the QIIME2 VSEARCH consensus classifier as the former classifiers did not provide species-level classification or could not be tuned to NanoCLUST data qualities (Murali *et al*., 2018; Bokulich *et al*., 2018). The VSEARCH classifier sets default identity (80%) and alignment length (80%) thresholds, ensuring that only high quality alignments are retained. Additionally, it collapses the top search hits by lowest common ancestor to ensure precision when closely related sequences exist in the reference database. We deviate in one other way from NanoCLUST: the minimum cluster size for HDBSCAN was recently decreased from 200 to 50 in NanoCLUST (as of February 15, 2021); BugSeq has yet to follow suit (as of March 5, 2021).

We evaluated our novel 16S analysis pipeline, BugSeq 16S, with publicly available full-length 16S sequencing data of known mock microbial communities. Cusco et al. sequenced the ZymoBIOMICS mock microbial community, containing eight bacteria in even abundance, in duplicate (Cuscó *et al*., 2019). Using NanoCLUST, there were 6 and 5 species correctly identified in the samples, with 4 and 2 false positive species, respectively (average precision 66%, average recall 69%). Using BugSeq 16S, we identify 7 bacterial species correctly in each sample, with 1 false positive in both (average precision 88%, average recall 88%). Specifically, NanoCLUST classified the *Listeria monocytogenes* sequences as *Listeria innocua*, the *Bacillus subtilis* sequences as *Bacillus halotolerans* and *Bacillus tequilensis*, and the *Escherichia coli* sequences as *Shigella flexneri*. BugSeq 16S identified the *Bacillus subtilis* sequences to the correct genus (without classifying to species level), and falsely identified an uncultured bacterium from the *Escherichia*/*Shigella* genus, in addition to the present *E. coli*. Revising the NanoCLUST minimum cluster size to 200 reads did not affect conclusions: there was still misidentification of the *B. subtilis* and *L. monocytogenes* reads, with *E. coli* no longer detected. On a more complex mock microbial community containing 20 bacteria spanning a 1000-fold concentration range (BEI HM-783D), NanoCLUST detected 5 species correctly with 8 false positive species; BugSeq 16S detected 6 species correctly with 1 false positive species (Cuscó *et al*., 2019). Recall was therefore 30% for BugSeq as compared to 25% for NanoCLUST, with precision of 85.7% for BugSeq and 38% for NanoCLUST. Full results are available in Table 1.

**Table 1:**
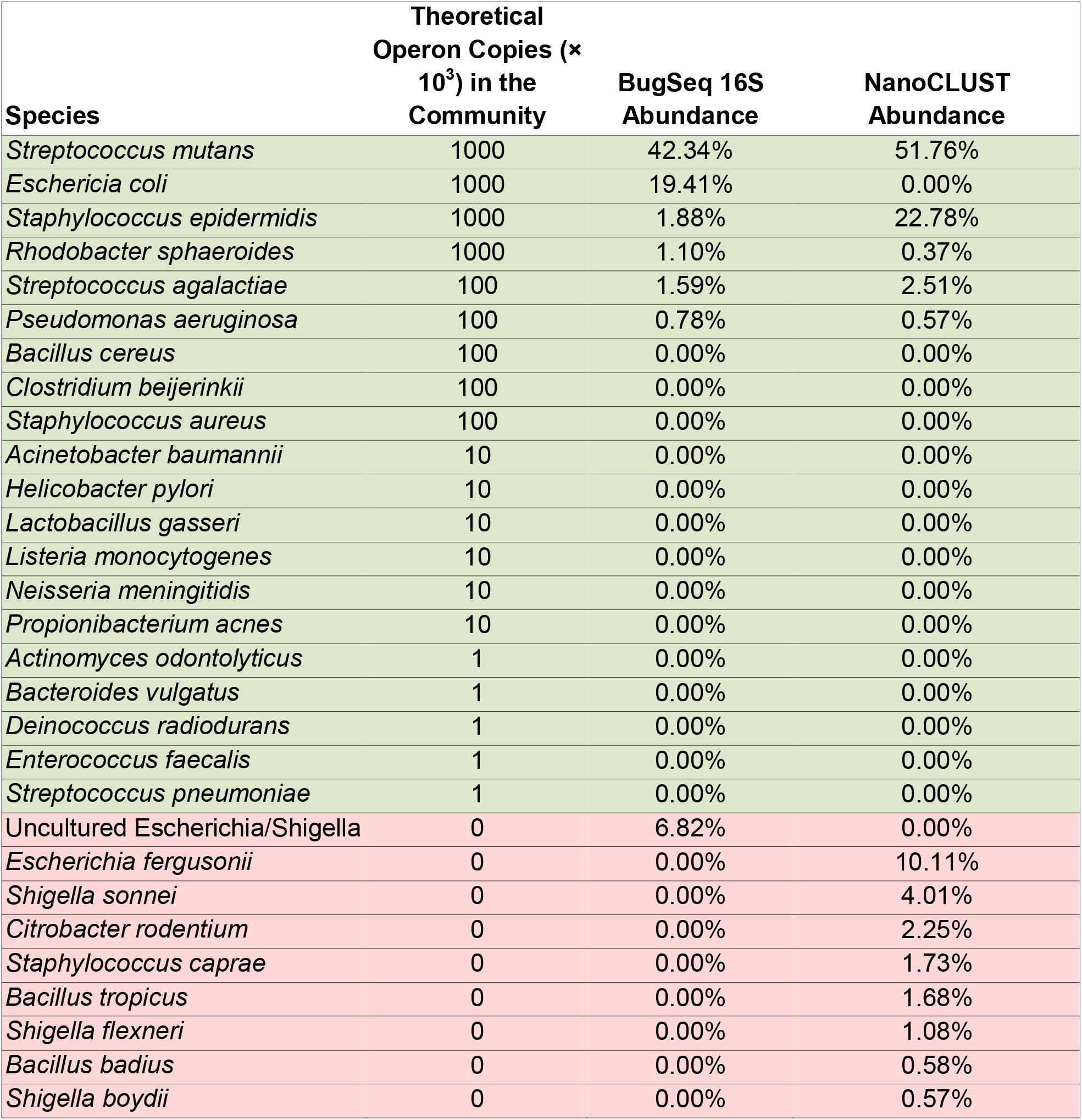
Species-level abundance in the HM-783D mock microbial community using BugSeq 16S and NanoCLUST. Green rows reflect species actually present in the sample, red rows reflect false positive species.

We next compared BugSeq 16S with NanoCLUST on publicly available full-length 16S sequencing data from patients with ventilator associated pneumonia (Maes *et al*., 2021). Twenty-nine bronchoalveolar lavage samples from 24 patients underwent microbial culture, multi-pathogen TaqMan array and nanopore 16S sequencing as previously described (Maes *et al*., 2021). We used a combination of culture and TaqMan array results as the gold standard. As TaqMan and culture results were censored by the original authors at a cycle threshold value of 32 and 10^4^ colony forming units per milliliter, respectively, only sensitivity of 16S sequencing could be calculated. Full data is available in the supplementary material.

At the species level, BugSeq 16S achieved better sensitivity (n=28/35, 80%) as compared to NanoCLUST (n=26/35, 74%). Specifically, patient 16 had an *E. coli* detected by culture and BugSeq 16S but NanoCLUST detected a mix of *Escherichia fergusonii, Shigella flexneri* and *Shigella sonnei*. Patient 17 has a *Serratia marcescens* detected by culture, TaqMan and BugSeq 16S, while NanoCLUST detected a *Serratia nematodiphila* and *Serratia surfactantfaciens*. BugSeq 16S achieved this greater species-level sensitivity while predicting fewer total species present in each sample. BugSeq 16S predicted a median of 13 (IQR=8) fewer species present in each sample as compared with NanoCLUST.

As BugSeq 16S collapsed results based on sequence similarity, it also correctly detected a *Stenotrophomonas spp*. in patient 1 (sample 2) which NanoCLUST mislabelled as *Stenotrophomonas pavanii* (ground truth: *Stenotrophomonas maltophilia*) and a *Haemophilus spp*. in patient 21 which NanoCLUST mislabelled as *Haemophilus parahaemolytics* (ground truth: *Haemophils influenzae*). Accounting for these confidence-aware classifications increases BugSeq 16S’s sensitivity to 85.7%.

In conclusion, BugSeq 16S builds on NanoCLUST to improve nanopore 16S sequencing analysis using a popular, highly accurate sequence classifier. This change results in significantly improved analysis accuracy on both mock microbial communities and real patient samples, and is free for academic use at <u>bugseq.com/free</u>.

## Notes

https://bugseq.com/free

### Competing Interest Statement

Ana Jung is an employee of BugSeq Bioinformatics Inc. Samuel Chorlton holds equity in BugSeq Bioinformatics Inc.

